# Biological networks with singular Jacobians: their origins and adaptation criteria

**DOI:** 10.1101/2021.03.01.433197

**Authors:** Tracey Oellerich, Maria Emelianenko, Lance Liotta, Robyn P. Araujo

## Abstract

This work is focused on Ordinary Differential Equations(ODE)-based models of biochemical systems that possess a singular Jacobian manifesting in non-hyperbolic equilibria. We show that there are several classes of systems that exhibit this behavior: a)systems with monomial-type interaction terms and b)systems with linear or nonlinear conservation laws. While models derived from mass-action principles often present with linear conservation laws stemming from the underlying biologic rationale, nonlinear conservation laws are more subtle and harder to detect. Nevertheless, in both situations the corresponding ODE system will contain non-hyperbolic equilibria. While having a potentially more complex dynamics and falling outside of the scope of existing theoretical frameworks, this class of systems can still exhibit adapting behavior associated with certain nodes and inputs. We derive a generalized adaptation condition that extends to singular systems and is compatible with both single-input/single-output and multiple-input/multiple-output settings. The approach explored herein, based on the notion of Moore-Penrose pseudoinverse, is tested on several synthetic systems that are shown to exhibit homeostatic behavior but are not covered by existing methods. These results highlight the role of the network structure and modeling assumptions when understanding system response to input and can be helpful in discovering intrinsic relationships between the nodes.

## 1. Introduction

Mathematically, singular Jacobians indicate a source of structural instability of the system and lead to non-hyperbolic equilibria which can change behavior, disappear or split into several equilibria as a result of a small perturbation. From biological standpoint, these systems are often excluded from consideration or, when possible, reduced to non-singular systems with more robust hyperbolic equilibria.

Our investigation is focused on two aspects related to these systems. First, we are interested in understanding the biological and modeling characteristics responsible for this behavior. Second, we study a subclass of singular problems that exhibit certain type of adaptation and develop the necessary theoretical tools for identifying this behavior.

An example of a well studied biological system with singular Jacobi matrix can be found in the supplement of Kholodenko et al.[13]. The system presented therein is part of the MAPK pathway and consists of 9 interacting molecules (*nodes*) with 3 different conservation laws. In this case, the system would have to be reduced by a dimension of at least 3 in order to eliminate singularities. Furthermore, the system contains Michaelis-Menten kinetics and reducing the system of this type can prove to be computationally demanding, especially as the dimension is increased. Systems such as the epidermal growth factor receptor (EGFR) pathway, which is known to exhibit adaptation [5, 9], can be modeled with a large number of nodes and contain multiple sources of singularities. There are other similar examples in the biological literature, all of which indicate the need for a more comprehensive analytical treatment, especially in relation to adaptation mechanisms. Elucidating fundamental properties of biological networks that are responsible for this function requires careful analysis of their structure.

Adaptation in the sense of asymptotic tracking of a ‘set-point’, has been widely explored in the literature [4, 12] at various levels ranging from the cell to the whole-organism level in mammals. At the cellular level, several types of adaptation have been studied in previous works, including perfect adaptation[9], fold-change detection (FCD)[19], absolute concentration robustness[18], homeostasis[20], and robust perfect adaptation[3]. All of these concepts share adapting behavior, although they highlight certain specific types of adaptive behavior. In [9], adaptation for cell signaling networks is defined as “a process where a system initially responds to a stimulus, but then returns to basal or near-basal levels of activity after some period of time”. Furthermore, for the adaptation to be considered perfect, the final level of response from the system must be the same as the level prior to any stimulus acting on the system ([14], [19], and [9]). In Muzzey et al.[16], for example, yeast osmoregulation analysis is performed to experimentally study the perfect adaptation exhibited by the system and the conditions under which is it observed.

Stated mathematically, perfect adaptation may be viewed as follows. Consider a certain dynamical system *ż* = *F* (*z, u*), where *z* = *z*(*t*) is an n-dimensional vector of state variables, *F* is the function describing system dynamics and *u* = *u*(*t*) is a generally time dependent input. Denote *y* = *h*(*z*) the corresponding output (“read-out” map). Let states, inputs, and outputs lie in particular subsets ℤ ⊂ ℝ^*n*^, 𝕌 ⊂ ℝ^*m*^, 𝕐 ⊂ ℝ^*q*^, respectively. Assume for each constant input *u*(*t*) = *ū*, there is a unique steady-state, denote it *σ*(*ū*), of the system (which depends on the particular input). This steady state is assumed to be globally asymptotically stable, although this assumption is not always apparent in the discussion. This system *perfectly adapts to constant inputs* provided that the steady-state output *h*(*σ*(*ū*)) equals some fixed *y*_0_ ϵ 𝕐, independently of the particular input value *ū* ϵ 𝕌. There are several reported instances of RPA occurring in biological systems, such as in bacterial chemotaxis ([21],[6], [2]), EGFR-regulated signaling pathways ([9],[17]), and transcription networks [10]. This notion is closely related to the phenomenon called *homeostasis* which is the biological property of maintaining a fixed state despite external perturbations, generally achieved by coupling a biological sensing mechanism to a feedback loop, so when perturbations attempt to change the state of the system, the feedback mechanism acts to resist these changes and restore the system to its default state. Under this definition one may distinguish between single-input/single-output and more general multi-input/multi-output systems. *Robust perfect adaptation (RPA)*, for instance, typically refers to the property of a biological system to return to the same activity level following any persistent change to the incoming signal received at the input node.

In any of the above interpretations, a biological network model encodes a set of interactions of a collection of molecules (*nodes*). These nodes could represent proteins, RNA transcripts, genes, or combinations. Nodes can encode and/or transmit a biochemical signal (e.g. proteins, RNA) and can undergo changes in their activity based on interactions with other nodes. In some systems, there will be only be one input node and one output node, while more general systems might exhibit more complex scenarios. In recent works, most notably in [3] and [20], mathematical conditions for adaptation were derived for single-input/single-output and multi-input/multi-output systems, respectively. These conditions were only applicable in the case of a hyperbolic steady state, ruling out potential singularities.

We will show that there is a possibility for singular systems to have nodes which exhibit adaptation despite not strictly satisfying the conditions for homeostasis or RPA derived in [3] and [20]. Furthermore, we derive a set of generalized conditions for adaptation suitable for this situation that are fully compatible with RPA and homeostasis frameworks and investigate the origins of singularities in these systems. We show that two important classes of systems exhibiting this behavior are the systems that include either a conservation law and/or nodes that model multimeric complexes rather than individual state variables. While representing the same biological mechanism, systems containing molecular complexes as individual nodes cannot be treated using standard RPA conditions. We give examples demonstrating that this more general definition of a node may produce RPA-capable network models with non-hyperbolic steady-states. We derive conditions that allow for the complete analysis of the RPA-capacity of such models.

In some cases, as we will show, it is possible to remove the singularity from the system. However, this requires knowing the system structure prior to any modifications. For systems with linear conservation laws, it may be easy to spot the cause of the singularity, but as can be seen in Mahdi et al.[15], identifying non-linear conservation laws within the system can be a strenuous and time consuming process. Furthermore, if the system is known to have several conservation laws or built-in assumptions affecting various nodes, removing the singularities may take longer than utilizing the general conditions for adaptation presented in this work. Having an alternate method for testing adaptation can eliminate the need to pre-process the system to remove singularities.

## 2. Mathematical Notations

Consider a system with state space denoted as *𝒫* ⊂ ℝ ^*N*^. The state space consists of *N* nodes, say **P**(*t*) = [*P*_1_(*t*), …, *P*_*N*_ (*t*)]^*T*^ ϵ *𝒫*, representing the interacting molecules of interest, such as proteins, RNA transcripts, genes.

Let *𝒰* ⊂ ℝ ^*M*^ be the *input* space and **U**(*t*) = [*U*_1_(*t*), *U*_2_(*t*), …, *U*_*M*_(*t*)]^*T*^ ϵ *𝒰* be the time dependent input to the system. For our systems, inputs are quantities which can be externally manipulated and applied. Examples include applying a drug to the system and increasing the dosage over time. The input could also be perturbations to the system coefficients used in order to monitor the effects on the system.

The *output* space will be *Y* ⊂ ℝ, and *h : 𝒫* → *Y*, 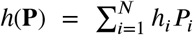 is the *output* which is intended to be kept constanby homeostasis. The output *h*(**P**) is a linear function of the state variables. By the choice of coefficients *h*_1_, we can choose the output to be a single state, or a weighted sum of multiple states in the system. Therefore, let *h* = [*h*_1_, …, *h*_*N*_] with 0 ≤ *h*_1_ 1 ≤ for all, be the (row) vector of coefficients which determine the weighted contribution from each node to the total output of the system.

Finally, let *F : 𝒫* × *𝒰* → ℝ^*N*^ be the continuously differentiable function describing the system *dynamics*. The biochemical system will describe the activities of the state variables over time as functions of the state variables and the inputs. The resulting dynamical system has the form:

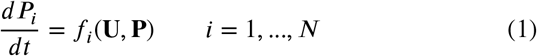

In matrix form, the nonlinear dynamical system (1) can be written as **P**^′^ = **F**(**P, U**).

The matrix *J* (**P, U**) = *D*_**P**_*F* (**P, U**) denotes the Jacobian of the system with respect to the system nodes, and the system **P**^f^ = *D*_**P**_*F* (**P**^*^, **U**)**P** represents the linearization of the system (1) at the steady state **P**^*^ = *g*(**U**).

Steady states of this dynamical system will satisfy

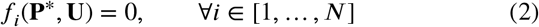

Most of the discussion in the literature has been focusing on steady states in which the eigenvalues of the Jacobian have non-zero real parts. The aim of this work is to understand the situations when a singular Jacobian at steady state might arise and to derive conditions for adaptation suitable for any type of steady state.

## 3. Origins of the singularities

First, let us discuss conditions under which singular systems may arise. In this section we will suppress dependence on input **U**, but the results hold true when the input-output map structure is imposed. We will also limit discussion to the system behavior at the network steady state.

### 3.1 Conservation laws

We propose (and prove) here that one possible reason for a singular Jacobian is the presence of a conservation law. For mass action systems, it is common to see a linear conservation law in place among 2 or more nodes within the system. The following result characterizes the Jacobian in this situation.

#### Proposition 3.1.

*Consider a biological system with N ≥* 2 *nodes which contains a linear conservation law. The system is defined as follows:*

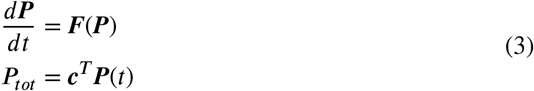

*where* ***c*** *≢* ***0*** *is a vector of coefficients in the conservation law and P*_*tot*_ *is the total concentration which remains constant. Then*, det(*J* (***P***(*t*))) = 0, ∀ *t ≥* 0.

The proof of Proposition 3.1 can be found in the Supplemental Notes S.1. The proof is a simple calculus exercise and relies on taking two derivatives of the conservation law, one with respect to time and the other with respect to a node, and seeing the effect on the structure of the Jacobian. In this case, it causes linear dependence amongst the rows corresponding to nodes in the linear conservation law. In particular, it is interesting to note this singularity is present for all time, meaning we do not need to evaluate at the network steady state to have a singular Jacobian.

The same cannot be said for a non-linear conservation law. First, we will consider a simplified version of the completely nonlinear case in which we will sum over nonlinear functions of one node.

#### Proposition 3.2.

*Consider a biological system with N ≥* 2 *nodes which contains a simple non-linear conservation law. The system is defined as follows:*

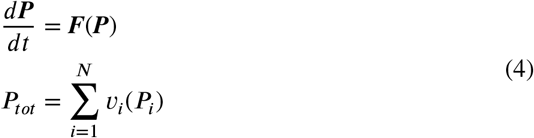

*where* ***v*** = *v*_1_(*P*_1_) … *v*_*n*_(*P*_*N*_) *≢* ***0***, *v*_i_ ϵC^2^(*𝒫*), *and for* 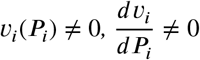 *and* 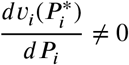 *centration which remains constant. (If we allow*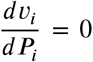, *this means that v*_*i*_(*P*_*i*_) = *b, where b is a constant. However, this would be independent of P*_*i*_.*) Then*, det(*J* (***P***^*^)) = 0 *for this system*.

The proof of Proposition 3.2 can be found in Supplemental Notes S.2. In this case, we do not see any singularities occurring unless we are at the network steady state. We can expand on this result by assuming we have a completely non-linear conservation law.

#### Proposition 3.3.

*Consider a biological system with N ≥* 2 *nodes which contains a nonlinear conservation law. Let* ***M*** = {*i*: *P*_*i*_ *appears in the conservation law*}, 2 ≤ *m* = | ***M*** | *≤ N. The system is defined as follows:*

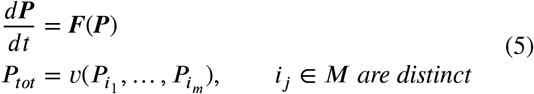

*where* 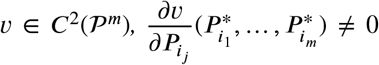 *for non-trivial steady state* ***P***^*^, *and P*_*tot*_ *is the total concentration which remains constant. Then*, det(*J* (***P***^*^)) = 0 *for this system*.

The proof of Proposition 3.3 can be found in Supplemental Notes S.3. To illustrate these results, consider the oxidation catalysis model (6) considered in Emelianenko et al. [8] which exhibits a nonlinear conservation law given by (7):

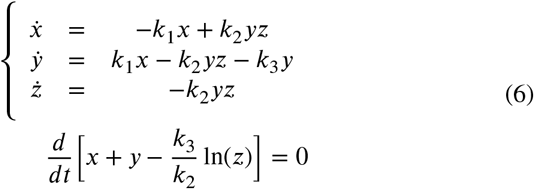

The system has the nontrivial steady state solution *x*^*^ = *y*^*^ = 0, *z*^*^ ≠ 0. The corresponding Jacobian evaluated at the steady state for this system is:

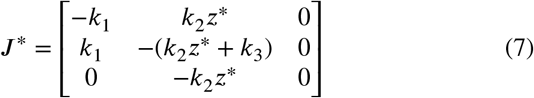

which clearly has det(*J* ^*^) = 0. This is a simple example of a singular system exhibiting a nonlinear conservation law. While it was relatively easy to detect conserved relation in this 3×3 system, more effort is necessary to do this in higher dimensional cases and the not all systems are amenable to existing techniques (see e.g. Mahdi et al.[15] and discussion therein).

### 3.2. Special network structures

While conservation laws contribute to the occurrence of singular Jacobians, they are not the only source. Another way for a system to exhibit a singularity at the steady state is if there is a multi-node complex located within the system which is modeled as a single node within the network, or if a certain type of interaction between nodes is giving rise to singularity. The below proposition details this case.

#### Proposition 3.4.

*Consider an N-node network, N ≥* 3, *in which one node, say P*_*N*_, *is a function of the other nodes within the network. Let the system be defined as:*

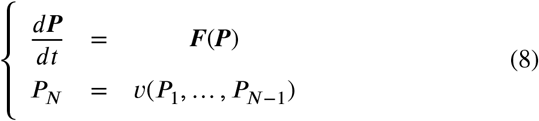

*where v* ϵ C^2^(*𝒫*^*N*−1^) *and at least one*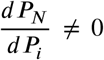 *for i* = 1, …, *N* − 1. *Then*, det(*J* (***P***^*^)) = 0 *for this system*.

The technical proof of this proposition is given in Supplemental Notes S.4. We provide examples realizing this scenario and its implications in Supplemental Notes S.5.

## 4. Adaptation in biological networks

Now that we have shown situations in which a singular Jacobian can occur, the next step is to examine how it affects the adaptation criteria defined in Araujo et al. [3]. Consider again the nonlinear dynamical system (1):

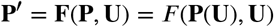

First, consider the full derivative of the system:

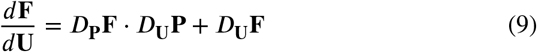

where *D*_**P**_**F** represents the Jacobian of **F** with respect to the nodes **P**, *D*_**U**_**P** represents the Jacobian of **P** with respect to the inputs **U**, and *D*_**U**_**F** represents the Jacobian of **F** with respect to the inputs **U**. Evaluating (9) at the steady state **P**^*^ = *g*(**U**) yields:

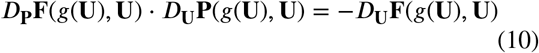

To simplify notation, all quantities will be evaluated at the steady state **P**^*^ = *g*(**U**) unless otherwise noted.

For all types of adaptation conditions, one needs to examine how the output node(s) react to changes in the input(s), i.e. understand the behavior of *D*_**U**_**P**, which leads to two cases:

1. det(*D*_**P**_**F**) ≠ 0 (non-singular, all eigenvalues are nonzero).
2. det(*D*_**P**_**F**) = 0 (singular, at least one eigenvalue is zero)

In what follows, we will outline the results available for Case 1 and extend them to Case 2.

### 4.1. Known results for non-singular systems and their consistency

Case 1 corresponding to det(*D*_**P**_**F**) ≠ 0 was examined in both [3] and [20] for the single-output and multiple-output cases, respectively. We will briefly discuss the correspondence between the two since it is instrumental for the extension considered below.

In the *multiple-output case*, [20] shows that a system will exhibit adaptation (homeostasis) if

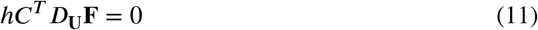

at steady state. Here *h* = [*h*_1_, …, *h*_*N*_] is the vector of coefficients which determine the weighted contribution from each node to the total output of the system and denotes the cofactor matrix of *D*_**P**_**F**.

It is not difficult to sketch the main ideas of the proof. Since *D*_**P**_**F** is assumed to be invertible, one can write the inverse as 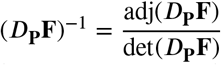, where adj(*D*_**P**_ **F**) is the transpose of the cofactor matrix. The cofactor matrix,, for *D*_**P**_**F** has entries defined as follows:

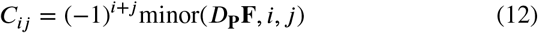

where the minor matrix is formed by removing the *i*-th row and the *j*-th column from *D*_**P**_**F**. Thus, as in [20], the problem becomes:

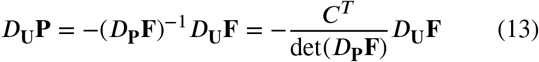

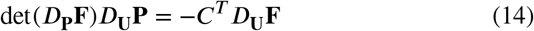

Now, we will examine the output of this system. To do so, we will isolate the effects of the inputs on the nodes which contribute to the overall output of the system by multiplying both sides by *h* = [*h*_1_ *h*_2_ … *h*_*N*_], which denotes the concentrations of each node in the output.

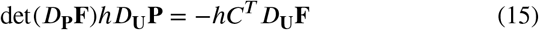

Finally, in order for the system to exhibit adaptation, the output of the nodes at steady state should be independent of the input, indicating that *hD*_**U**_**P** = 0. Thus, at an attracting steady state *g*(**U**), we must have

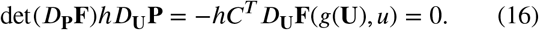

This is precisely the homeostasis condition given in [20].

In the *case of a single input/single output* system, we will consider a single input, **U**(*t*) = *I*(*t*) (we will also use *I* when referring to the index of the node the input is being applied to), and output, 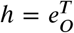, where *O* is the index of the output node. A biochemical system for which det(*D*_**P**_**F**) ≠ 0 at the system’s steady state will exhibit RPA-type adaptation if, at steady state, det(*M* _*IO*_) = 0, where *M* _*IO*_ is the minor matrix of *D*_**P**_**F** obtained by removing row *I* and column *O*, as shown in [3]. This is consistent with the result (16) above. Indeed, let *P*_*I*_ be the input node for the system and *P*_*O*_ denote the output node. Thus, the problem becomes:

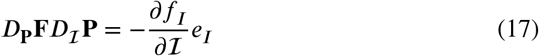

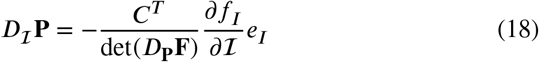

For RPA to occur, the output node must always returns to the same activity level, indicating that 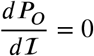. We can isolate the output node equation in the system as follows:

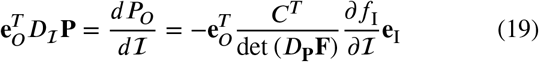

where 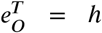 in the homeostatic setting. However,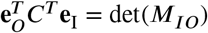 where *M*_*IO*_ defined as before. Thus

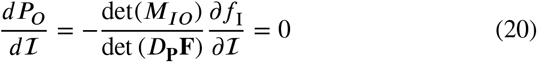

There are two possibilities: det(*M_IO_*) = 0 or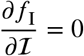. The latter is addressed as a special case of the system having a “paradoxical component” (such as in a bifunctional enzyme [11]) by Araujo and Liotta in the supplemental notes of [3]. In the absence of this behavior, det(*M*_*IO*_) = 0.

Thus, for a system to exhibit RPA, we have the following two conditions:

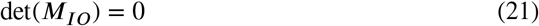

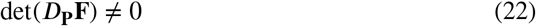

where (21-22) is precisely the *RPA equation* and *RPA constraint* as defined in [3].

This demonstrates that both conditions fall under the same umbrella. By removing the assumption of invertibility, we will now give a natural extension of both conditions.

## 5. Generalized adaptation criterion for singular systems

Let us consider the second case, det(*D*_**P**_**F**) = 0. In this case, *D*_**P**_**F** is not invertible. Therefore, we need to use other means in order to isolate *D*_**U**_**P**. We use singular value decomposition of the Jacobian to derive the following result.

### Proposition 5.1.

*A biochemical system* (1) *with inputs* ***U*** ϵ ℝ^*M*^*for which* det(*D*_***P***_***F***)(*P* ^*^) = 0 *exhibits adaptation if*

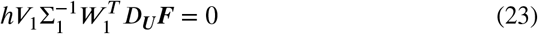

*at the system’s steady state P* ^*^, *where V*_1_, Σ_1_, *and W*_1_ *are the components of the singular value decomposition of D*_***P***_***F***.

*Proof*. Consider the singular value decomposition (SVD) of *D*_**P**_**F**, where *D*_**P**_**F** is of rank *r* < *N*:

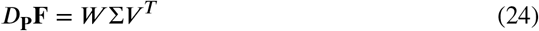

where *W, V*, Σ ϵ ℝ^*N*×*N*^, *W* is the matrix containing the left singular vectors, *V* is the matrix containing the right singular vectors, and Σ is a diagonal matrix containing the singular values in decreasing order. This decomposition exists for any matrix regardless of the nature of its eigenvalues [7]. Since *D*_**P**_**F** is of rank *r* < *N*, we can use the reduced form of the SVD:

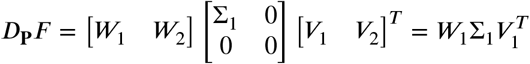

where Σ_1_ ϵ ℝ^*r*×*r*^ is the diagonal matrix of non-zero singular values of *D*_**P**_**F**, *W*_1_ ϵ ℝ^*N*×*r*^ is the orthogonal matrix of left singular vectors associated with nonzero singular values, *W*_2_ ϵ ℝ^*N*×(*N*−*r*)^ is the orthogonal matrix of left singular vectors associated with 0 singular values, *V*_1_ ϵ ℝ^*N*×*r*^ is the orthogonal matrix of right singular vectors associated with nonzero singular values, and *V*_2_ ϵ ℝ^*N*×(*N*−*r*)^ is the orthogonal matrix of right singular vectors associated with 0 singular values. The Moore-Penrose pseudoinverse of *D*_**P**_**F**, denoted by †, is given by 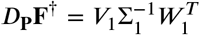. We can now simplify the problem as follows:

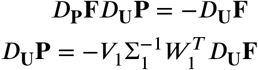

Now, we will examine the output of this system by isolating the effects of the inputs on the output nodes as done in the previous case:

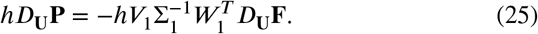

In order for the system to exhibit adaptation, the output at steady state should be independent of the input, indicating that *hD*_**U**_**P** = 0. Conclusion follows.□

Now as a special case consider single-input/single-output scenario, i.e. *M*= 1 and *h*_*i*_ = 0, ∀*i* ≠ *O*.

**Corollary 5.1**. *Let be the input node for the system and PO denote the output node. A biochemical system* (1) *for which* det(*D*_**P**_***F***)(*P*^*^) = 0 *exhibits RPA if, at steady state*,

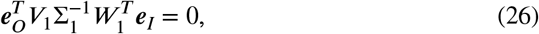

*where V*_1_, *W*_1_, *and* Σ_1_ *are obtained from the singular value decomposition of* det(*D*_***P***_***F***).

The result follows directly from Proposition 5.1 by fixing 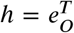 and 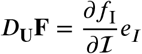:

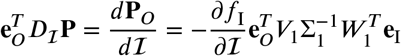

where *V*_1_, *W*_1_ ϵ ℝ^*N*×*r*^, Σ_1_ ϵ ℝ^*r*×*r*^ are obtained from the SVD of *D*_**P**_**F** which has rank *r < N*.

We have now shown how to extend adaptation results to systems with singular Jacobians. The practical utility of theoretical results is summarized in the flowchart presented in Figure 1. We provide a simple numerical demonstration of this approach for the case consistent with singularity case described in Section 3.2 in Supplemental Notes S.5.

**Figure 1:**
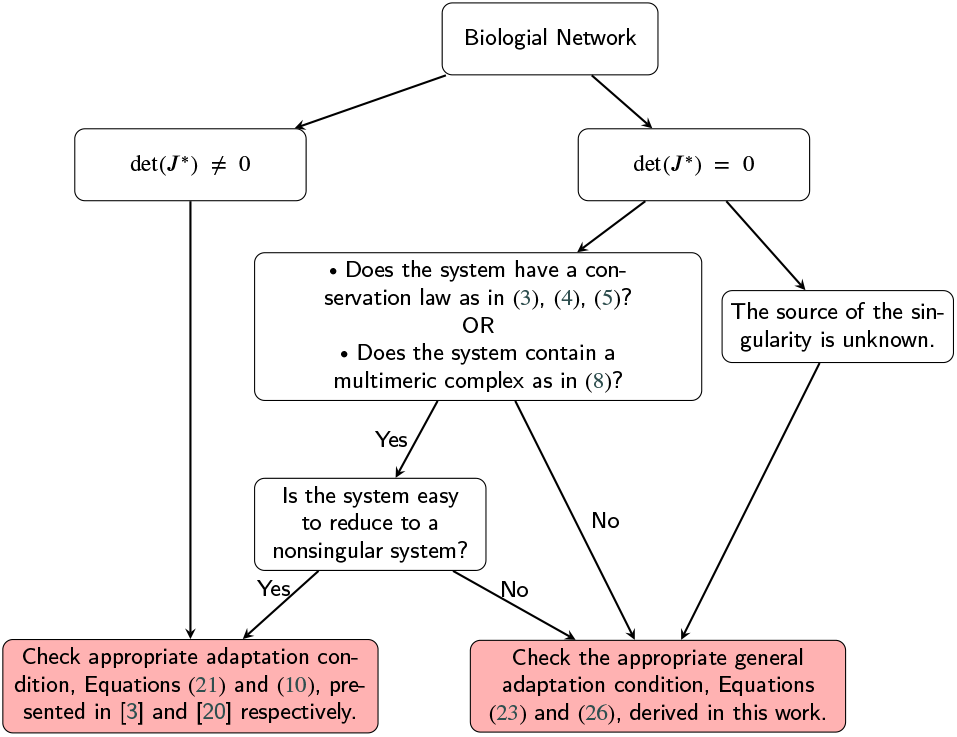
Flowchart detailing process of checking adaptation.

## 6. Discussion

In this work, we have examined several types of biochemical reaction networks which give rise to a singular Jacobian, including systems with a special type of network structure and those containing a linear or nonlinear conservation law. We compared and contrasted different types of adaptative behaviors governing evolution of living organisms, including single-input/single-output and multiple-input/multiple-output scenarios. Following the dynamical systems formalism we separated these systems into classes based on the singular or nonsingular behavior of their Jacobians evaluated at the steady state. It’s important to note that all analytical approaches to the study of RPA to date, to our knowledge, are predicated on the existence of hyperbolic equilibria. But as we show here, mathematical models of biological signaling networks readily give rise to singular Jacobian matrices in a variety of circumstances, and thus cannot be analysed for their adaptation potential by existing methods. Examples of biological systems and numerical validation of the new approach are provided to demonstrate practical importance of this class of problems and the feasibility of the singular behavior that cannot be analyzed using standard RPA conditions.

Putting adaptation criteria aside, there are a few potential applications of the analytical results connecting singularities with certain types of system characteristics. First, from the results presented herein, we can conclude that the presence of a singularity can indicate existence of the underlying conservation laws or flaws in certain modeling assumptions. If discovered, these features can significantly reduce problem dimension and ultimately remove the singular behavior leading to a more robust and stable mathematical formulation. This may be particularly useful in the context of data-driven modeling of biological cascades that amplify inaccuracies in model formulations yet can be dramatically improved through the process outlined above. In such a modeling cycle, there is a possibility of a spurious singularity created for instance by a suboptimal choice of kinetic constants, or vice versa, an inherently singular system containing a conservation law could lose that structure for the same reason. Having additional knowledge provided in this investigation can help guide this process towards biologically meaningful formulations.

The question of which network structures give rise to singularities needs to be explored further. While a definitive answer was obtained in the case of conservation laws and multimeric complexes/interactions of general type that are modeled as individual nodes, there may still be others. It also remains to be shown that if a system with a singularity exhibits a generalized RPA, the corresponding reduced system will necessarily exhibit a regular RPA behavior. The numerical example involving a monomial complex provided in the Supplemental Notes indicates this might be true in this specific case, but further work is needed to make a more general statement. The modified RPA condition can also be used to measure the amount of perturbation necessary to destroy adaptation, which can lead to a quantifiable prediction of the role each node plays in the system response to external stimuli. For singular systems with no known underlying structure, this condition allows to investigate adaptive behavior for any given node or a combination of nodes. For singular systems with existing conservation laws, it allows to directly test for adaptation while avoiding computationally intensive algebraic manipulations.

Furthermore, one may ask a question of how the adaptation mechanism react to changes in the network structure, such as permuting the system topology as well as input/output structures, similar to the work performed in [1], [14], and [20]. We conjecture that adapting behavior is a fragile property of a living system that may be easily lost by relatively simple network modifications. This will be the subject of future investigations.

## 7. Acknowledgements

The authors gratefully acknowledge fruitful discussions with Alessandra Luchini and Abdulaziz Alaraini. TO was partially funded by the George Mason University Provost PhD award under the Industrial Immersion Program. RPA is the recipient of an Australian Research Council (ARC) Future Fellowship (project number FT190100645) funded by the Australian Government.

## Supplemental Notes

### S.1 Proof of Proposition 3.1

*Proof*. Consider a biological system with *N* ≥ 23 nodes which contains a linear conservation law defined as follows:

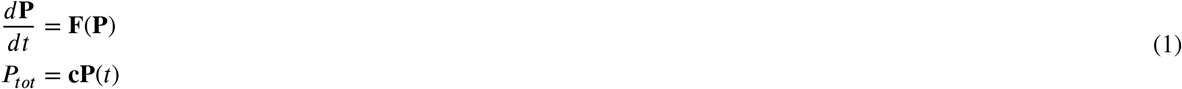

where **c** ≢**0** is a vector of constants and *P*_*tot*_ is the total concentration which remains constant. Let **F** = [*f*_1_ … *f*_*N*_]^*T*^ and **P** = [*P*_1_ … *P*_*N*_]^*T*^.

If we take the derivative with respect to time of the conservation law, we have:

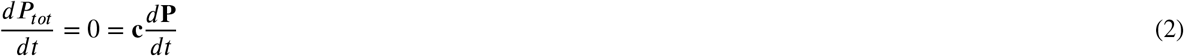

Therefore, we can solve for one 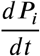, say we solve for 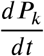, where *c*_*k*_ ≠0. Since **c** ≢0, we have:

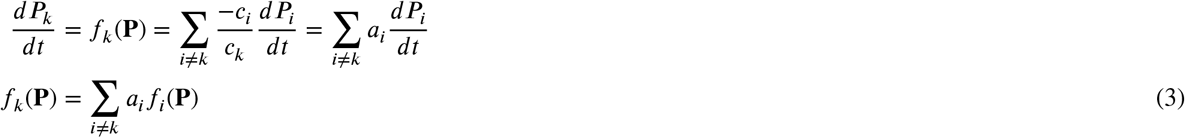

where 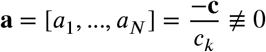.

Now we will look at he Jacobian of the system **F** with respect to the nodes **P**. This is given by:

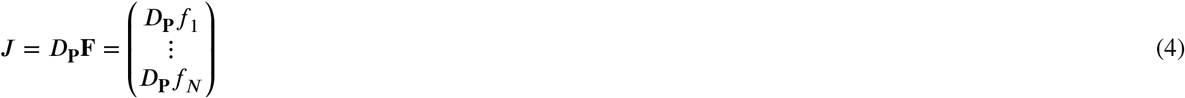

Where

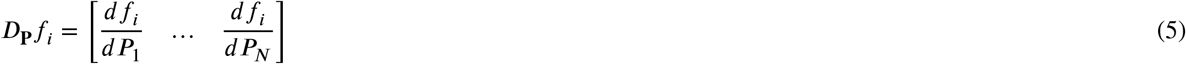

Now, looking at *D*_**P**_*f*_*k*_, we see this row vector is:

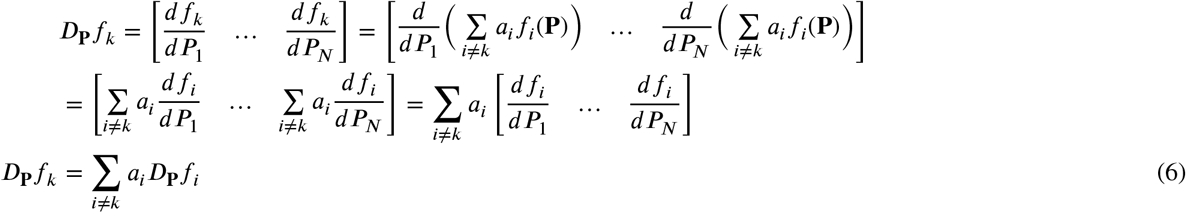

Therefore, the k-th row of the Jacobian, *D*_**P**_*f*_*k*_, can be written as a linear combination of the other rows of the Jacobian. Thus, the Jacobian is singular and the determinant is det(*J*) = 0. □

### S.2 Proof of Proposition 3.2

*Proof*. Consider a biological system with *N* ≥ 2 nodes which contains a simple non-linear conservation law. The system is defined as follows:

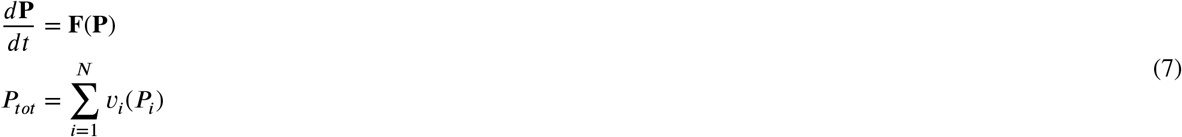

where **v** = [*v*_1_(*P*_1_) … *v* _*N*_ (*P*_*N*_ ≢ **0**, each *v* _*i*_(*P*_*i*_) ϵ C^2^, for *v* _*i*_(*P*_*i*_) ≠ 0 we have 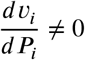 and 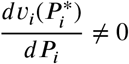, and *P*_*tot*_ is the total concentration which remains constant.

If we take the derivative with respect to time of the conservation law, we have:

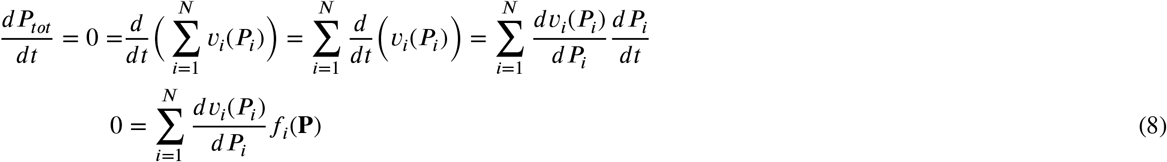

Now we will solve for one 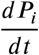, say we solve for 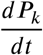, where 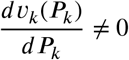. Then we have:

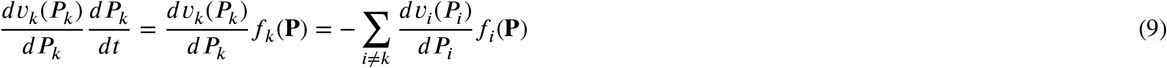

The Jacobian for this system will is defined as:

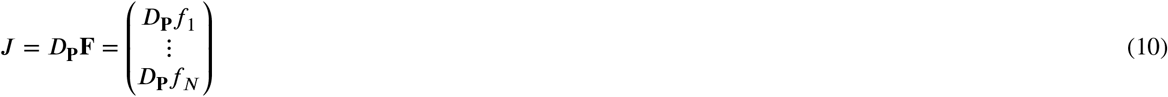

where

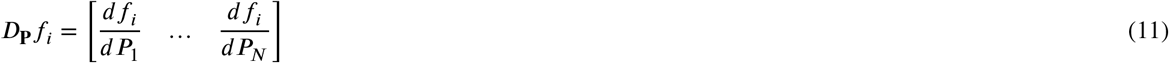

Note for *f*_*k*_, the corresponding row in the Jacobian, *D*_**P**_*f*_*k*_ will be:

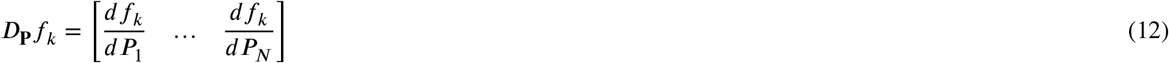

Now we will find the entries for this row:

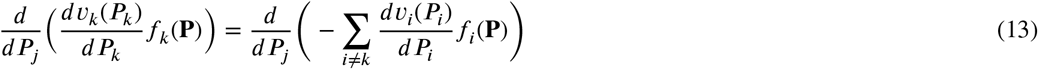

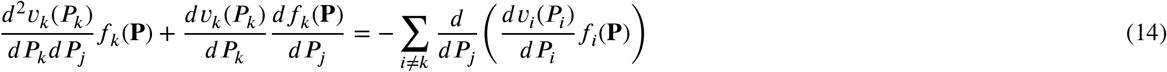

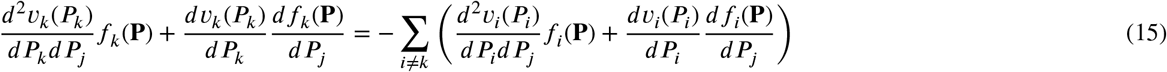

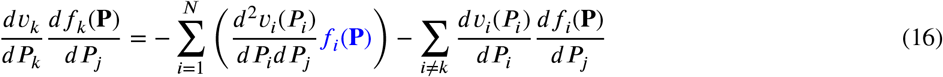

In this setting, the Jacobian is certainly not guaranteed to be singular since the 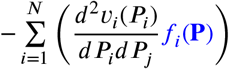 term in each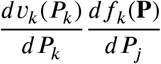 is unlikely to be equal to 0 for all *j*.

So suppose we evaluate the Jacobian at the network steady state, denoted 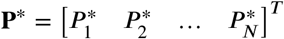. The steady state will satisfy

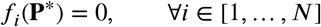

Now, in this case, we have the Jacobian:

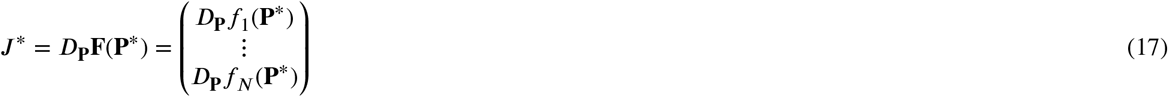

Where

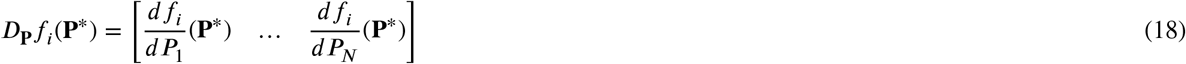

From before, we have:

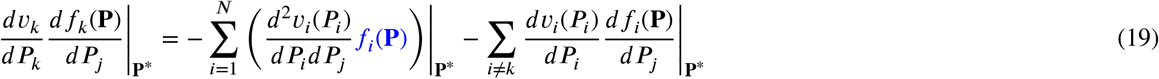

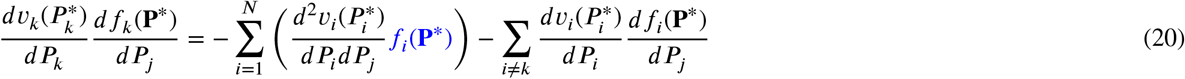

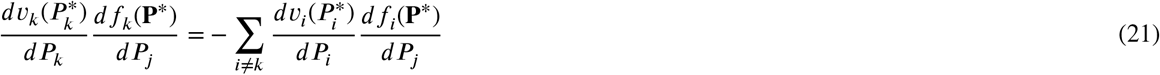

Since we defined 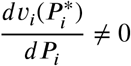 when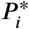 is the network steady state solution, we can invert 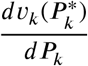 and therefore have:

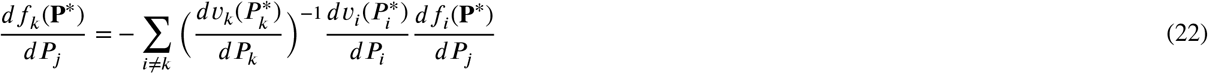

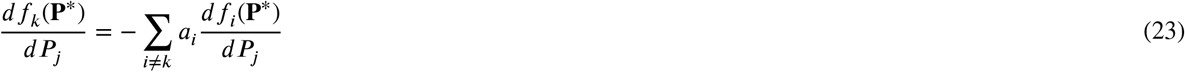

where each 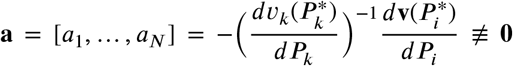. Now for *f*_*k*_, the corresponding row in the Jacobian when evaluated at the network steady state, *D*_**P**_*f*_*k*_(**P** *) will be:

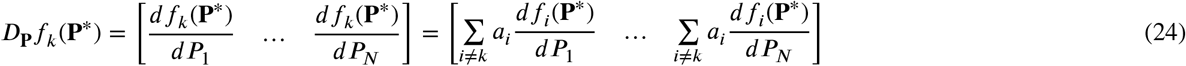

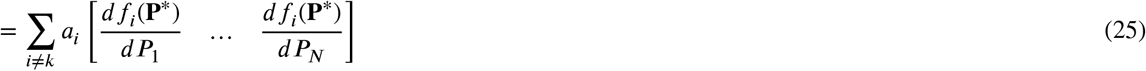

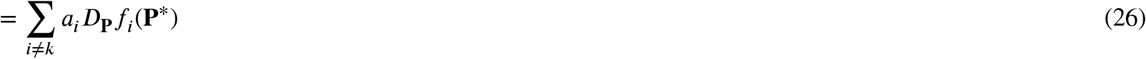

Therefore, the k-th row of the Jacobian evaluated at the network steady state is a linear combination of the other rows of the Jacobian. Thus, the determinant of the Jacobian at steady state is det(*J*\*) = 0 and the matrix is singular. □

### S.3 Proof of Proposition 3.3

*Proof*. Consider a biological system with *N* ≥ 2 nodes which contains a nonlinear conservation law. Let *M* = {*i*: *P*_*i*_ appears in the conservation law}, 2 ≤ |*M*| ≤ *N*. The system is defined as follows:

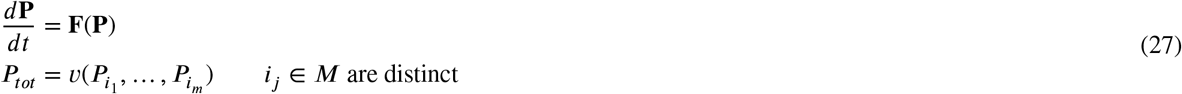

where 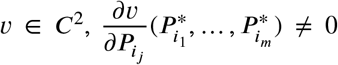 for non-trivial steady state **P**\*, and *P*_*tot*_ is the total concentration which remains constant. Let **F** = [*f*_1_ … *f*_*N*_]^*T*^ and **P** =[*P*_1_ … *P*_*N*_]^*T*^.

If we take the derivative with respect to time of the conservation law, we have:

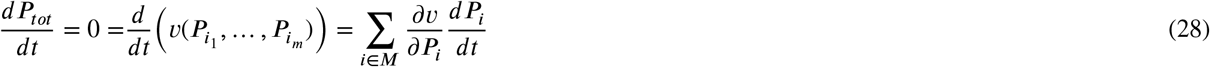

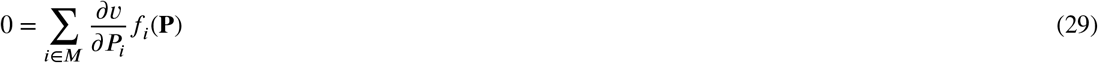

Now we will solve for one 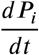, say we solve for 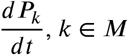, *k* ϵ *M*. Then we have:

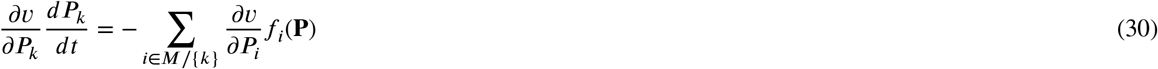

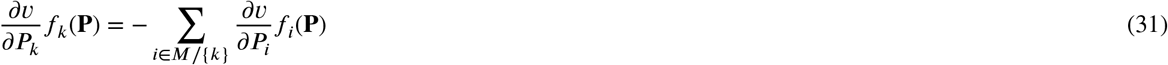

The Jacobian for this system will be defined as:

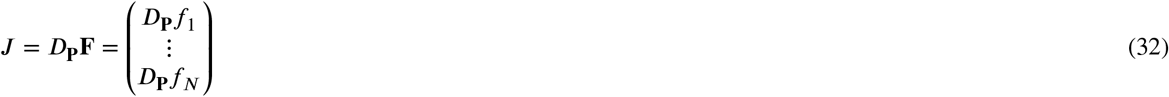

where

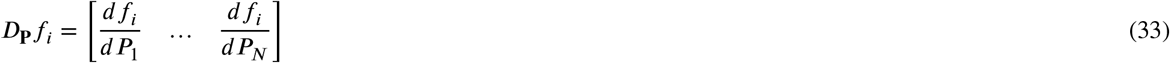

Note for *f*_*k*_, the corresponding row in the Jacobian, *D*_**P**_*f*_*m*_ will be:

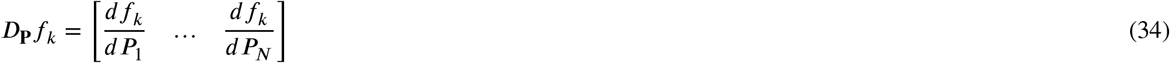

Now we will find the entries of this row.

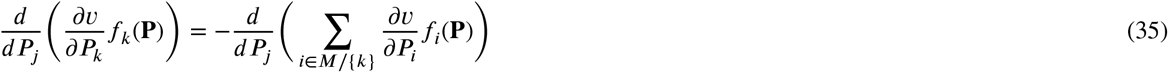

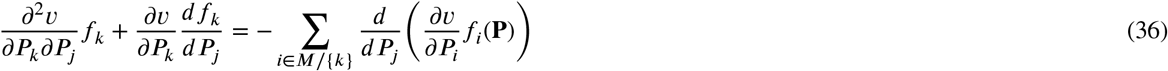

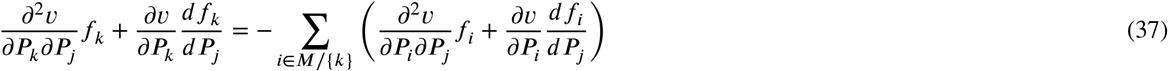

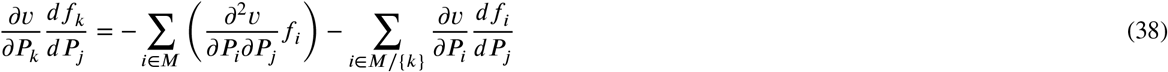

In this setting, the Jacobian is not guaranteed to be singular since to the 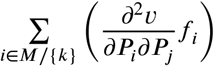 term in each 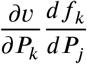 is not guaranteed to be 0 for all *j*.

Now suppose we evaluate the Jacobian at the network steady state, denoted 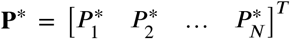. The steady state will satisfy

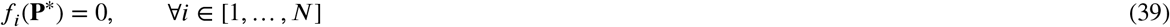

Now, in this case, we have the Jacobian:

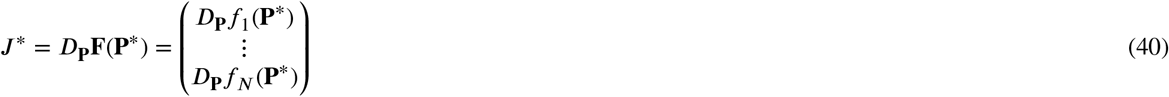

where

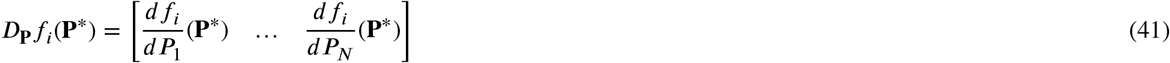

From before, we have:

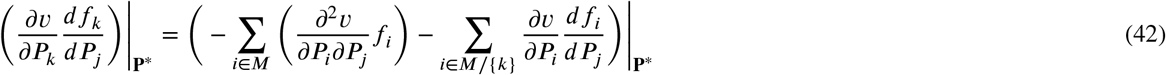

**Figure 1:**
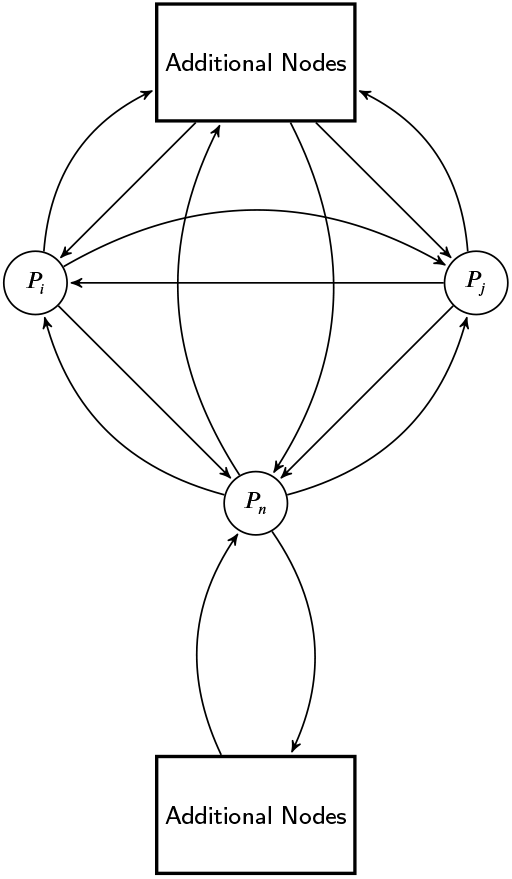
Illustration of a network considered in Proposition 3.4.

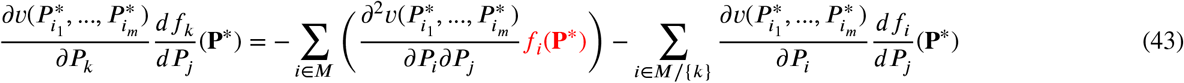

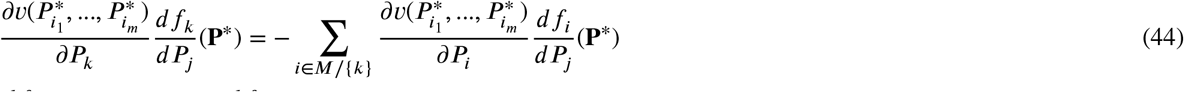

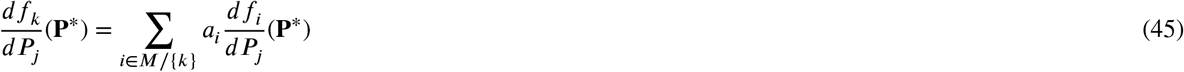

where each 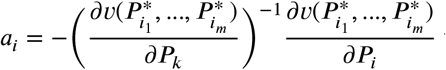 will be a nonzero constant.

Now for *f*_*k*_, the corresponding row in the Jacobian when evaluated at the network steady state, *D*_**p**_ *f*_*k*_(**P**\*) will be:

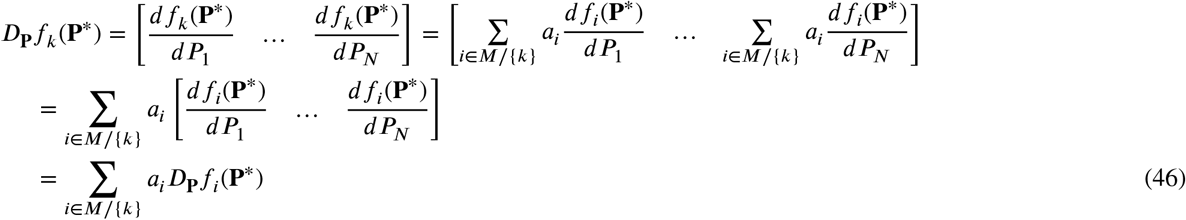

Therefore, the k-th row of the Jacobian evaluated the the network steady state is a linear combination of *m* − 1 rows. Thus, the determinant of the Jacobian at steady state is det(*J*\*) = 0 and the matrix is singular. □

### S.4 Proof of Proposition 3.4

*Proof*. Consider a system of *N* nodes, *N* ≥ 3, in which one node, say *P*_*N*_, is a function of the other nodes within the network, as depicted in Figure 1. Let the system be defined as:

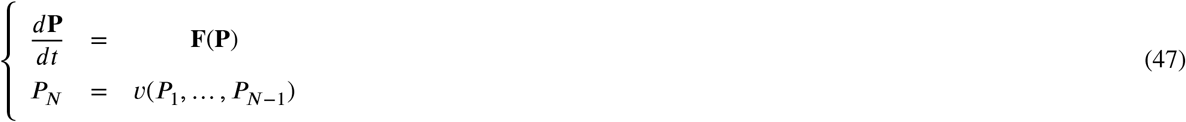

where at least one 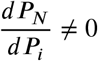 for *i* = 1, …, *N* − 1. Since we know *P*_*N*_ = (*P*_1_, …, *P*_*N*−1_), we have:

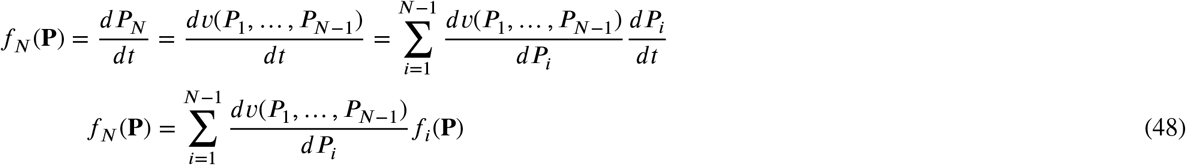

Then the Jacobian of this system, *D*_**P**_(**F**) is:

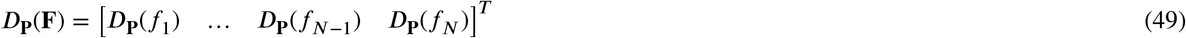

Where

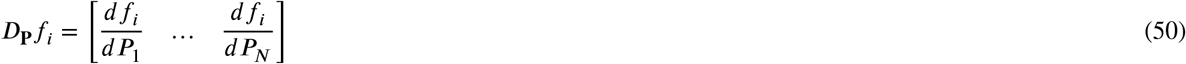

At the steady state *P*\*,

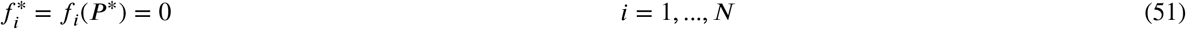

Now, we will consider the *N*-th row of the Jacobian in particular. The terms of this row are given by:

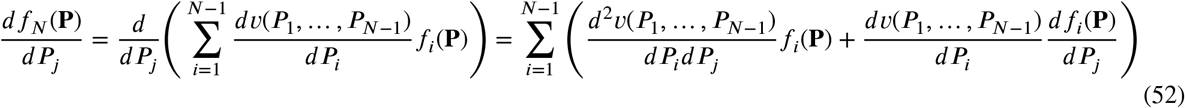

Evaluating at the network steady state, we have:

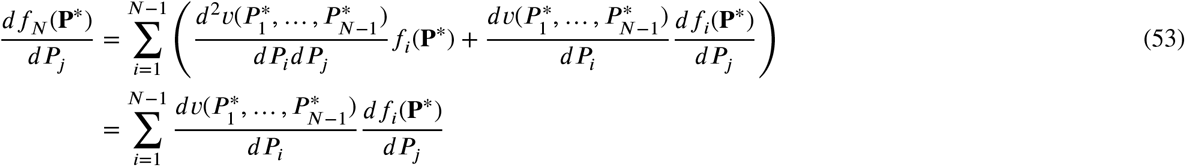

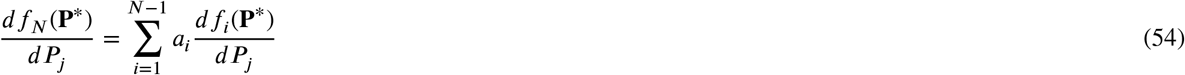

Where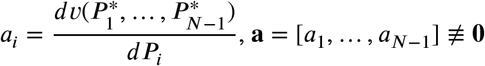.

Therefore, the *N*-th row of the Jacobian evaluated at the network steady state is:

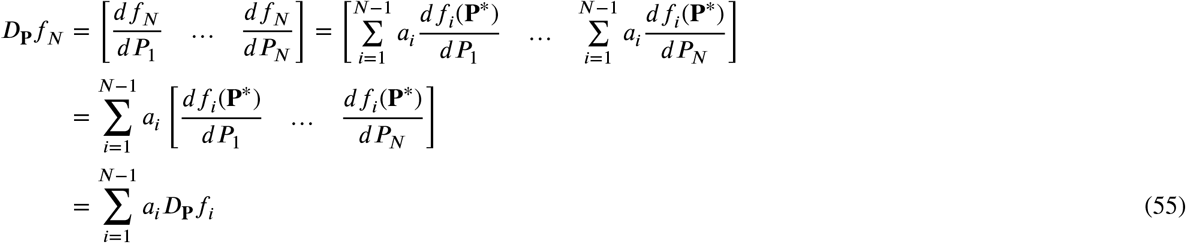

Therefore, the rows of *D*_**P**_**F**(**P**) are not linearly independent since the last row in a linear combination of previous rows. Thus det(*D*_**P**_**F**(**P**)) = 0 and the Jacobian evaluated at the steady state is singular. □

### S.5 Numerical example

We have performed analysis of several models to validate the newly derived adaptation conditions.

System (56) consisting of five nodes including a compound *P* = *CD* was considered in [1] and is represented schematically in Figure 2.

**Figure 2:**
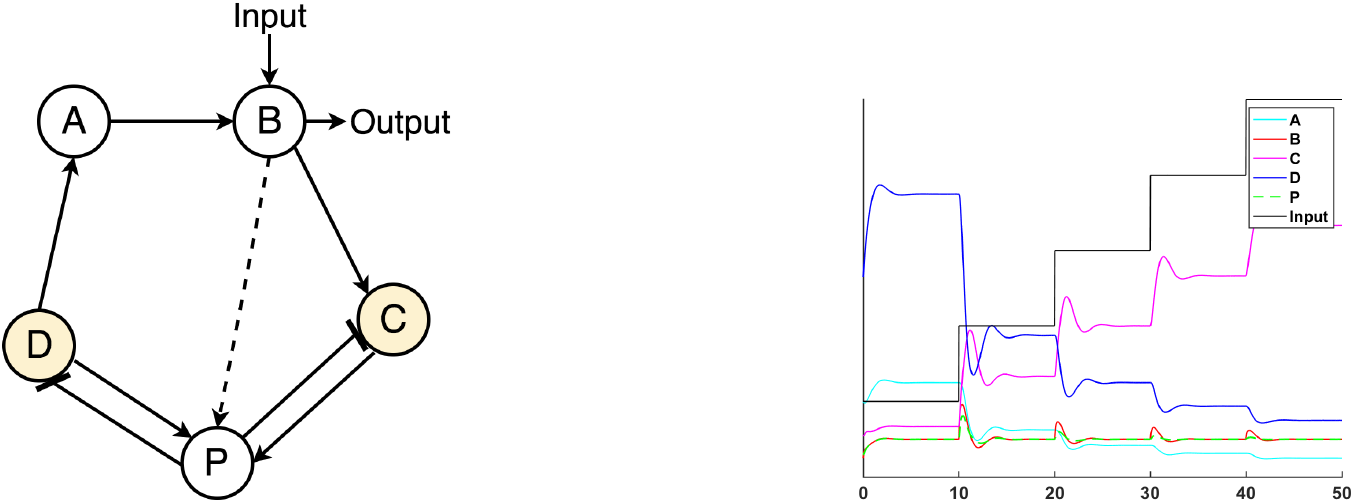
(left) Schematic of the singular system (56) presented in Araujo et al [1], which exhibits RPA and satisfies the conditions for RPA in the presence of a singular Jacobian. For this system, the input and output node are the same. (right) The output for each node over time when using a step function input. Node *B* exhibits the RPA property. Furthermore, node *P* also exhibits RPA.

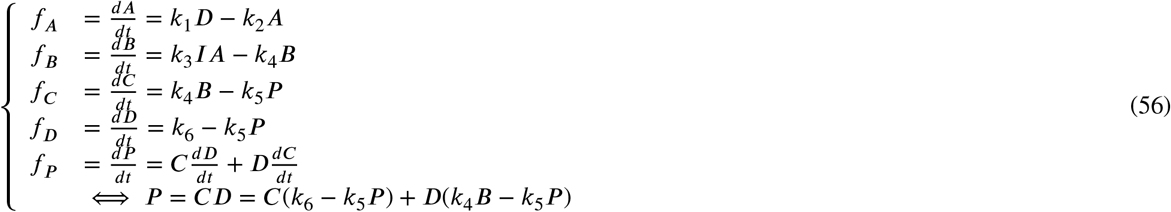

For this example, at steady state, we have

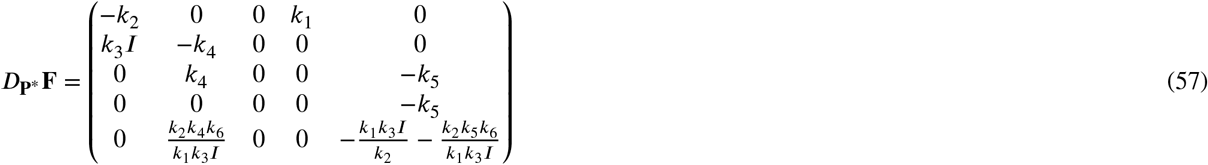

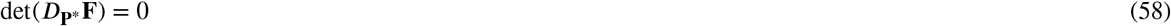

Therefore, it does not satisfy the standard RPA conditions of case 1 and we must look at the extended condition presented Corollary 5.1 (singular case). Once the reduced SVD is completed, we see that 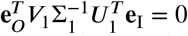 at steady state, indicating that this system exhibits RPA by Corollary 5.1. Indeed, we may observe this by numerically solving the system for certain randomly chosen coefficients and initial states, as demonstrated in Figure 2.

Note that this system is a version of the model presented in Ferrell [2]. The original system consisted of 4 nodes and a compound of two nodes (see Figure 3). For the 5-node system, the compound was treated as a separate node and added to the system. The original system is shown below:

**Figure 3:**
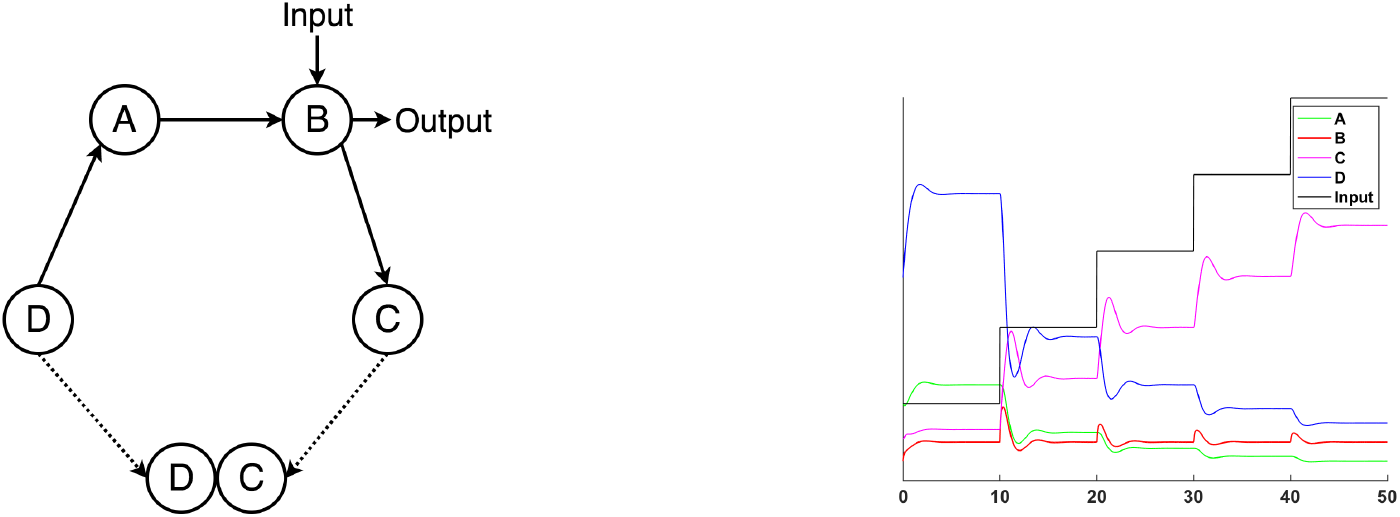
(left) Schematic of system (59) presented in Ferrell [2], which exhibits RPA and satisfies the RPA constraint and the RPA condition presented in Araujo et al. [1]. This system was adapted by [1] to be a 5 node system by treating the term *C* · *D* as a new node. (right) The output for each node over time when using a step function input. We can see that node *B* exhibits the RPA property.

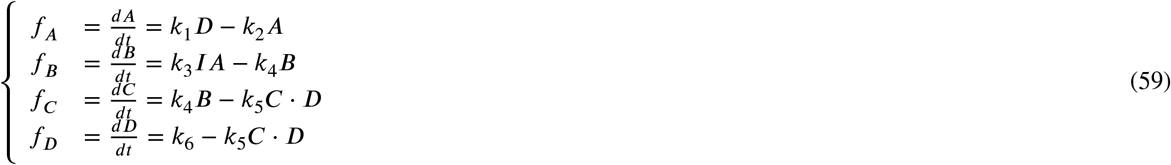

For the 4 node system at steady state, we have:

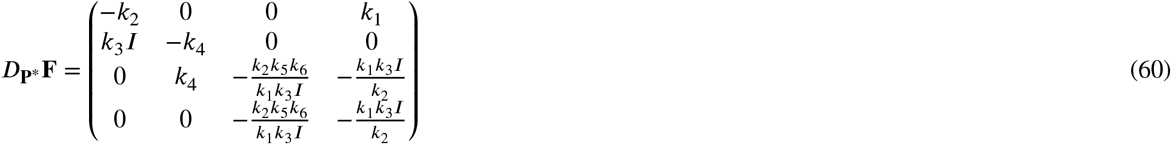

However, in this system, we have:

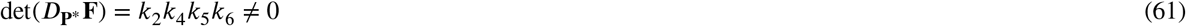

and therefore, the 4-node system is non-singular and belongs to case 1. Calculating det(*M*_*IO*_) at steady state for this system, where *B* is treated as the input and output node, yields:

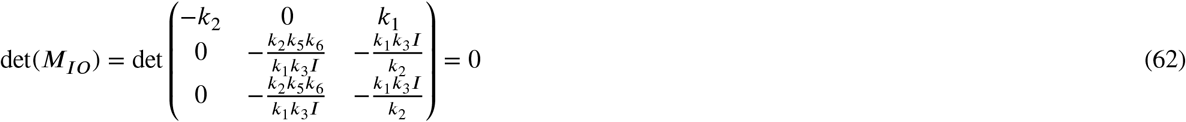

Therefore the four node system exhibits RPA. This can also be seen numerically as shown in Figure 3.

The examples above demonstrate that *modeling the biochemical compounds in different ways may have an impact on the analysis* due to the emergence of the singular behavior predicted by Proposition 3.4. Due to the versatile nature of the extended adaptation condition derived in the main paper, both situations can be successfully analyzed and same conclusion is reached in both cases.

